# Biologically plausible phosphene simulation for the differentiable optimization of visual cortical prostheses

**DOI:** 10.1101/2022.12.23.521749

**Authors:** Maureen van der Grinten, Jaap de Ruyter van Steveninck, Antonio Lozano, Laura Pijnacker, Bodo Rückauer, Pieter Roelfsema, Marcel van Gerven, Richard van Wezel, Umut Güçlü, Yağmur Güçlütürk

## Abstract

Blindness affects millions of people around the world, and is expected to become increasingly prevalent in the years to come. For some blind individuals, a promising solution to restore a form of vision are cortical visual prostheses, which convert camera input to electrical stimulation of the cortex to bypass part of the impaired visual system. Due to the constrained number of electrodes that can be implanted, the artificially induced visual percept (a pattern of localized light flashes, or ‘phosphenes’) is of limited resolution, and a great portion of the field’s research attention is devoted to optimizing the efficacy, efficiency, and practical usefulness of the encoding of visual information. A commonly exploited method is the non-invasive functional evaluation in sighted subjects or with computational models by making use of simulated prosthetic vision (SPV) pipelines. Although the SPV literature has provided us with some fundamental insights, an important drawback that researchers and clinicians may encounter is the lack of realism in the simulation of cortical prosthetic vision, which limits the validity for real-life applications. Moreover, none of the existing simulators address the specific practical requirements for the electrical stimulation parameters. In this study, we developed a PyTorch-based, fast and fully differentiable phosphene simulator. Our simulator transforms specific electrode stimulation patterns into biologically plausible representations of the artificial visual percepts that the prosthesis wearer is expected to see. The simulator integrates a wide range of both classical and recent clinical results with neurophysiological evidence in humans and non-human primates. The implemented pipeline includes a model of the retinotopic organisation and cortical magnification of the visual cortex. Moreover, the quantitative effect of stimulation strength, duration, and frequency on phosphene size and brightness as well as the temporal characteristics of phosphenes are incorporated in the simulator. Our results demonstrate the suitability of the simulator for both computational applications such as end-to-end deep learning-based prosthetic vision optimization as well as behavioural experiments. The modular approach of our work makes it ideal for further integrating new insights in artificial vision as well as for hypothesis testing. In summary, we present an open-source, fully differentiable, biologically plausible phosphene simulator as a tool for computational, clinical and behavioural neuroscientists working on visual neuroprosthetics.

## Introduction

Globally, as per 2020, an estimated 43.3 million people were blind (1). For some cases of blindness, visual prosthetics may provide a promising solution. These devices aim to restore a rudimentary form of vision by interacting with the visual system using electrical stimulation (2–4). In particular, our work concerns prosthetic devices that target the primary visual cortex (V1). Despite recent advances in the field, more research is required before cortical prosthesis will become clinically available. Besides research into the improvement of the safety and durability of cortical implants (5, 6), a great portion of the research attention is devoted to optimizing the efficacy, efficiency, and practical usefulness of the prosthetic percepts. The artificially induced visual percepts consist of patterns of localized light flashes (‘phosphenes’) with a limited resolution. To achieve a functional level of vision, scene-processing is required to condense complex visual information from the surroundings in an intelligible pattern of phosphenes (7–13). Many studies employ a simulated prosthetic vision (SPV) paradigm to non-invasively evaluate the functional quality of the prosthetic vision with the help of sighted subjects (9, 14–16) or through ‘end-to-end’ approaches, using in silico models (7, 12, 17). Although the aforementioned SPV literature has provided us with some fundamental insights, an important drawback is the lack of realism and biological plausibility of the simulations. At this stage of the development, given the steadily expanding empirical literature on cortically-induced phosphene vision, it is both feasible and desirable to have a more phenomenologically accurate model of cortical prosthetic vision. Such an accurate simulator has already been developed for retinal prostheses (15), which has formed an inspiration for our work on simulation of cortical prosthetic vision. Thus, in this current work, we propose a realistic, biophysically-grounded computational model for the simulation of cortical prosthetic vision. Our simulator integrates empirical findings and quantitative models from the literature on cortical stimulation in V1. The elements that are modeled in our simulator include cortical magnification, current-dependent spread of activation and charge-dependent activation thresholds. Furthermore, our simulator models the effects of specific stimulation parameters, accounting for temporal dynamics. A schematic overview of the pipeline and some example outputs are displayed in Figure 1. Our simulator runs in real-time, is opensource, and makes use of fully differentiable functions which allow for gradient-based optimization of phosphene encoding models. This design enables both simulations with sighted participants, as well as end-to-end optimization in machinelearning frameworks, thus fitting the needs of fundamental, clinical and computational scientists working on neuroprosthetic vision.

**Fig. 1.**
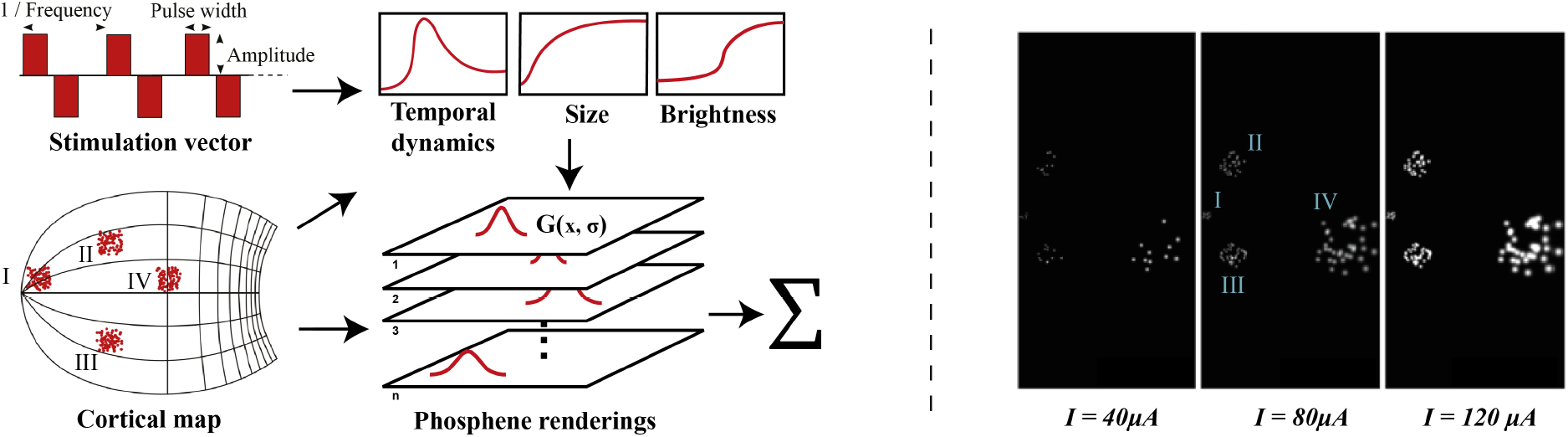
Left: schematic illustration of our simulator pipeline. Our simulator is initialized with electrode locations on a visuotopic map of the visual cortex. Each frame, the simulator takes a set of stimulation parameters that for each electrode specify the amplitude, pulse width, and frequency of electrical stimulation. Based on the electrode locations on the cortical map and the stimulation parameters, the phosphene characteristics are estimated and for each phosphene the effects are rendered on a map of the visual field. Finally, the phosphene renderings are summed to obtain the resulting simulated prosthetic percept. Right: Example renderings after initializing the simulator with four 10 × 10 electrode arrays (size: 5 × 5 millimetres) placed in the left hemisphere. The output is visualized for 166*ms* pulse trains with stimulation amplitudes of 40, 80, and 120*μA*, a pulse width of 170*μs*, and a frequency of 300*Hz*. In these example frames, we can observe the effects of cortical magnification, thresholds for activation, current-dependent spread (size) and proportion (brightness) of cortical tissue activation.

### A. Background and related work

#### A.1. Cortical prostheses

Early attempts by Brindley and Lewin, and Dobelle successfully reported to reliably induce the perception of phosphenes (described as localized round flashes of light) via electrical stimulation of the cortical surface (18, 19). More recent pre-clinical studies demonstrate promising results concerning the safety and efficacy of long-term stimulation in the primary visual cortex, either via surface electrodes (20, 21) or with intracortical electrodes (6, 22, 23). Other studies that performed V1 stimulation in sighted subjects (24, 25) and nonhuman primates (5, 26) have shown similar success. Some milestones include the implantation of over 1000 electrodes in a monkey’s visual cortex (5), and the testing of a preliminary artificial vision system that presents visual information from the surroundings to a blind subject using a penetrating multi-electrode array in the visual cortex (6). Taken together, the previous literature provides strong evidence for the clinical potential of cortical prostheses for the blind.

#### A.2. Perceptual reports on cortical prosthetic vision

In our simulator, we integrate empirical findings and quantitative models from existing literature on electrical stimulation in the visual cortex. Stimulation in V1 with intracortical electrodes is estimated to activate tens to thousands of neurons (27), resulting in the perception of often ‘featureless’ white dots of light with a circular shape (6, 18, 20, 22, 23). Due to the cortical magnification (the foveal information is represented by a relatively large surface area in visual cortex), the size of the phosphene is strongly correlated with its eccentricity (24, 25). Furthermore, phosphene size, stimulation threshold (defined as the minimum current to reliably produce a visible phosphene) and brightness are reported to be dependent on stimulation parameters such as the pulse width, train length, amplitude and frequency of stimulation (6, 20, 22–25). To account for these effects, we integrated and adapted previously proposed quantitative models that estimate the charge-dependent activation level of cortical tissue (24, 25, 28–31). Furthermore, our simulator includes a model of the temporal dynamics, observed by (23), accounting for response-attenuation after prolonged or repeated stimulation, as well as the delayed ‘offset’ of phosphene perception.

#### A.3. Simulated prosthetic vision

A wide range of previous studies has employed SPV with sighted subjects to non-invasively investigate the usefulness of prosthetic vision in everyday tasks, such as mobility (13, 14, 16, 32, 33), hand-eye coordination (34), reading (16, 35) or face recognition (36, 37). Several studies have examined the effect of the number of phosphenes, spacing between phosphenes and the visual angle over which the phosphenes are spread (e.g., (14, 34, 38–40)). The results of these studies vary widely, which could be explained by the difference in the implemented tasks, or, more importantly, by the differences in the simulation of phosphene vision. Most of the aforementioned studies used highly simplified phosphene simulations, with equally-sized phosphenes that were uniformly distributed over the visual field (informally referred to as the ‘scoreboard model’). Furthermore, most studies assumed either full control over phosphene brightness or used quantized levels of brightness (e.g. “on” / ”off”), but did not provide a description of the associated electrical stimulation parameters. Even studies that have explicitly made steps towards more realistic phosphene simulations, by taking into account cortical magnification or using visuotopic maps (34, 41, 42), did not model temporal dynamics or provide a description of the parameters used for electrical stimulation. Some recent studies developed descriptive models of the phosphene size or brightness as a function of the stimulation parameters (24, 25). Another very recent study has developed a deep-learning based model for predicting a realistic phosphene percept for single stimulating electrodes (43). While these studies have made important contributions to improve our understanding of the effects of different stimulation parameters, they do not provide a full simulation model that can be used for the functional evaluation of cortical visual prosthetic systems. This is what we aim to achieve in the current study. Meanwhile, a realistic and biologically-plausible simulator has already been developed for retinal prosthetic vision (Pulse2Percept, (15)), which takes into account the axonal spread of activation along ganglion cells and temporal nonlinearities to construct plausible simulations of stimulation patterns. Even though scientists increasingly realize that more realistic models of phosphene vision are required to narrow the gap between simulations and reality (13, 44), a biophysically-grounded simulation model for the functional evaluation of cortical prosthetic vision remains to be developed. Realistic SPV can aid technological developments by allowing neuroscientists, clinicians and engineers to test the perceptual effects of changes in stimulation protocols, and subsequently select stimulation parameters that yield the desired phosphene percepts without the need for extensive testing in blind volunteers. A realistic simulator could also be used as support in the rehabilitation process, assisting clinicians and caregivers in identifying potential problematic situations and adapt preprocessing or stimulation protocols accordingly (44).

#### A.4. Deep learning-based optimization of prosthetic vision

SPV is often employed to develop, optimize and test encoding strategies for capturing our complex visual surroundings in an informative phosphene representation. Numerous scene-processing methods have been proposed in the literature, ranging from basic edge detection or contour-detection algorithms (45–47) to more intelligent deep learning-based approaches, which can be tailored to meet task-specific demands (7, 8, 10, 13, 36, 48, 49). The advantage of deep learning-based methods is clear: more intelligent and flexible extraction of useful information in camera input leads to less noise or unimportant information in the low-resolution phosphene representation, allowing for more successful completion of tasks. Some recent studies demonstrated that the simulation of prosthetic vision can even be incorporated directly into the deep learning-based optimization pipeline for end-to-end optimization (7, 12, 17). With end-to-end optimization, the image processing can be tailored to an individual user or specific tasks. Here, the usefulness of the prosthetic vision is evaluated by a computational agent, or decoder model, which assists in the direct optimization of the stimulation parameters required to optimally encode the present image. Note that one requirement of the optimization pipeline is that the simulator makes use of fully differentiable operations to convert the stimulation parameters to an image of phosphenes. In the current study, we have adapted and improved upon the framework of (7) to demonstrate the compatibility of our realistic simulator for end-to-end optimization. The currently proposed simulator can handle temporal sequences and can be compared to the aforementioned work; our experiments explore a more biologically grounded simulation of phosphene size and locations. Furthermore, instead of a more abstract or qualitative description of the required stimulation (‘on’ / ‘off’), we included a biophysical model for predicting the perceptual effects of different stimulation parameters such as the current amplitude, the duration, the pulse width and the frequency. This opens new doors for optimization of the stimulation parameters in realistic ranges: although technological developments advance the state-of-the-art hardware capabilities rapidly, cortical prosthesis devices will be operating under energy constraints, due to both hardware limitations as well as safety limits regarding neurostimulation (50, 51). Deep learning methods trained in tandem with a biologically plausible phosphene simulator can be leveraged to produce constrained optimal stimulation paradigms that take into account these limitations, allowing for safe and viable stimulation protocols to be developed.

## Methods

Our simulator is implemented in Python, using the PyTorch deep learning library (52). The simulator makes use of differentiable functions which, given the entire set of phosphenes and their modelled properties, calculate the brightness of each pixel in the output image in parallel. This architecture makes our model memory intensive, but allows for fast computations that can be executed on a GPU. Each frame, the simulator maps electrical stimulation parameters (stimulation current, pulse width and frequency) to a predicted phosphene perception, taking into account the stimulation history. In the sections below, we discuss the different components of the simulator model, followed by a description of some showcase experiments. Our simulator can be imported as a python package and the source code is available on GitHub ^1^.

### B. Visuotopic mapping

Our simulator can be flexibly initialized with different modes for the phosphene topological mapping. In the most basic mode, a custom list of phosphene locations within the visual field can be provided by the user, expressed in polar coordinates. Alternatively, the phosphene locations can be calculated from a list of electrode locations on a flattened cortical map of V1. For mapping phosphene locations from the cortical electrode locations to the neuroprosthesis user’s visual field, our simulator uses the reverse wedge-dipole visuotopic model of V1, proposed by (53). This model maps a complex polar coordinate *z* = *re^i,θ^* in the visual field, to a cortical location *w* in one hemisphere of V1, following the visuotopic relationship

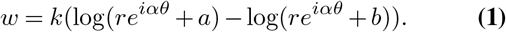

Here *r* and 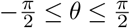 are the eccentricity and azimuth of the point in the visual field, the parameter *α* controls the shear of the ‘wedge map’, *k* is a scaling factor that scales the mapping to realistic proportions in cortical distance (millimeters), and *a* and *b* are parameters that control the singularities of the dipole model. For the mapping from cortical coordinates to phosphene location, we use the inverse of equation 1, which is given by

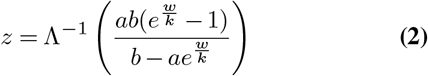

for the inverse shearing operation

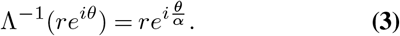

The visuotopic model also provides us with the cortical magnification *M*, which defines the relative amount of cortical tissue that is involved in processing of visual information, depending on the eccentricity in the visual field. The cortical magnification is given by the derivative of equation 1 along the horizontal meridian:

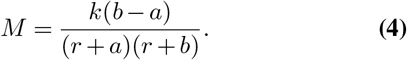

Here, *M* is given in millimetres of cortical surface per degree of visual angle. The parameters of the models are configurable. The default values are specified in section G. Note that in our simulation software, we provide the option of substituting equations 1–4, with other estimates described such as the mono- or dipole model in (53, 54).

### C. Phosphene size

Based on a model by (28), the phosphene size (in degrees),

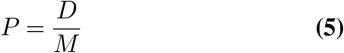

is obtained via an estimation of the current spread from the stimulating electrodes, where

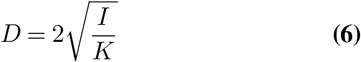

is the diameter of the activated cortical tissue (in *mm*), for stimulation current *I* (in *μA*) and excitability constant *K* (in *μA/mm*^2^). Note that the cortical magnification factor M is obtained in equation 4. The default value for *K* is specified in section G. In our simulation software, we provide the option to substitute equation 6 with an estimate by (25). Based on verbal descriptions (6, 22, 23), phosphenes are shown as Gaussian blobs with two standard deviations set equal to the phosphene size *P*, such that 95% of the Gaussian falls within the predicted phosphene size.

### D. Phosphene brightness

In our simulator, the brightness and detection threshold of each phosphene are based on a model of the intracortical tissue activation in response to electrical stimulation with biphasic square pulse trains. Our model assumes brightness and detection thresholds of phosphene perception to be primarily correlated with the deposited charge, and accounts for the relative inefficiency of higher stimulation frequencies, longer pulse-widths, and longer train durations, as found in (6, 20, 24). Our simulator models the combined effects of these stimulation parameters as follows: First, we subtract from the stimulation amplitude *I*_stim_ a leak current *I*_0_, which represents the ineffective component of the stimulation input, and a memory trace *Q* (further explained in section F) that accounts for the decreased neural excitability after prior stimulation. *I*_0_ is set equal to the rheobase current (the absolute threshold for continuous stimulation at infinite duration), following prior literature on the strength-duration relationship of neural tissue activation for isolated single-pulse trials (29). To calculate the effective stimulation current of trains of pulses, the remaining current amplitude is multiplied with the duty cycle of the stimulation signal (*Pw* · *f*, the fraction of one period in which the signal is active), such that

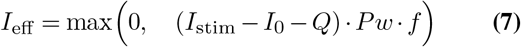

for pulse width *Pw* and frequency *f*. Then, the cortical tissue activation is estimated by integrating the effective input current over multiple frames, using a leaky integrator model. For each frame with duration Δ*t*, the estimated cortical activation is updated as

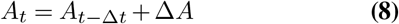

with

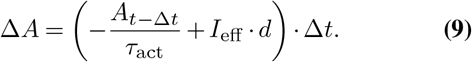

Here, *τ*_act_ is the time constant of the activation decay in seconds and *d* ∈ (0, 1] is a parameter that scales the duration of the stimulation relative to the frame duration. By default, *d* is set to 1 to simulate a stimulation duration equal to the frame duration, where the total pulse train duration is controlled with the number of successive frames in which stimulation is provided to the simulator. Finally, if the cortical activation reaches the detection threshold (explained in section E), the phosphene is activated with a brightness equal to the sigmoidal activation

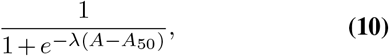

where *λ* is the slope of the sigmoidal curve and *A*_50_ is the value of *A* for which the phosphene reaches half the maximum brightness.

### E. Stimulation threshold

Our simulator uses a thresholding model based on psychometric data from (6). Phosphenes are only generated when the cortical tissue activation (explained in section D) reaches the activation threshold *A*_thr_, which is obtained for each electrode separately upon initialization of the simulator. To introduce a degree of variability between electrodes, *A*_thr_ is sampled from the normal distribution

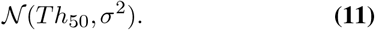

The default values of the 50% probability threshold *Th*_50_, and the standard deviation *σ* are fit on data from (6) and can be found in section G. Note that, by default, the detection thresholds remain constant after initialization. However, in accordance to the user requirements, the values can be flexibly adjusted or re-initialized manually.

### F. Temporal dynamics

Using a memory trace of the stimulation history, our simulator accounts for basic accommodation effects of brightness for prolonged or repeated stimulation, as described in prior work (23). Each frame, the memory trace is dynamically updated as follows:

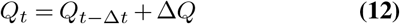

with

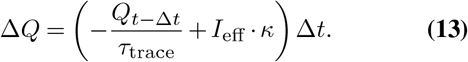

Here, *τ*_trace_ is the time constant of the trace decay in seconds, and the parameter *κ* controls the input effect. Note that the memory trace is used for the phosphene brightness and not the phosphene size. As there is little experimental data on the temporal dynamics of phosphene size in relation to the accumulated charge, only the instantaneous current is used in the calculation of the phosphene size.

### G. Parameter estimates

By default, our model uses the parameters specified below. Unless stated otherwise, these parameter estimates were obtained by fitting our model to experimental data using the SciPy Python package, version 1.9.0 (55). More details on the comparison between the models’ predictions and the experimental data can be found in the next section. Note that the parameter settings may strongly depend on the specific experimental conditions (such as the type of electrodes).

- In equations 1, 2, 3, 4 we use *a* = 0.75, *k* = 17.3, *b* = 120, and *α* = 0.95, based on a fit by (53) on data of the human V1 from (56).
- In equation 6, the parameter *K* is set to 675*μA/mm*^2^, following an estimate by (57), who measured the responses to intracortical stimulation in V1 at different current levels.
- In equation 7, we use a rheobase current *I*_0_ = 23.9*μA* based on a fit on data from (6). Here, we used the strength-duration curve for tissue-excitability *Q*_thr_ = *I*_0_ (*c* +*t*), with minimal input charge *Q*_thr_, chronaxie parameter *c* and total stimulation duration *t* as described in (29).
- In equation 9, the parameter *d* is set equal to 1. The parameter *τ*_act_ is set equal to 0.111*s* to reflect qualitative descriptions found in (23). Note: this parameter is not obtained by fitting to experimental data.
- In equation 10, we use a slope *λ* = 19.2 · 10^7^ and offset *A*_50_ = 1.06 · 10^−6^ for the brightness curve, based on a fit of our model on data by (6).
- The descriptive parameters of the distribution 11, are set to *Th*_50_ = 9.14 · 10^−8^ and *σ* = 6.72 · 10^−8^, based on a fit on psychometric data by (6).
- In equations 12 and 13, we use *τ*_trace_ = 1.97 * 10^3^*s* and *κ* = 14.0. These values are based on a fit of our model to data from (23).

## Experiments and Results

In this section we present computational experiments and comparisons with the literature to validate the performance, biological realism, and practical usability of our simulator.

### H. Performance

We tested the computational efficiency of our simulator, by converting a pre-processed example video^2^ (1504 frames) into simulated phosphene images, for different numbers of phosphenes, and at varying image resolutions. The simulator was run on a CUDA-enabled graphics card (NVIDIA© A30) and each setting was run five times. The results are displayed in Figure 2. The lowest measured frame rate (10.000 phosphenes at a resolution of 256 × 256) was 28.7 frames per second. Note that the missing combinations in Figure 2 indicate that the required memory exceeded the capacity of our GPU, as the simulation of large numbers of phosphenes at high resolutions can be memory intensive.

**Fig. 2.**
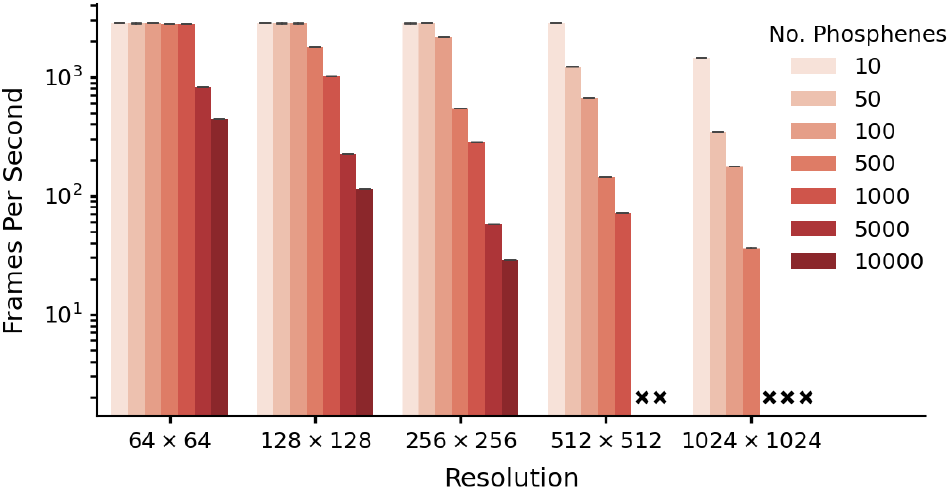
Performance as a function of resolution and number of phosphenes. The data is based on 5 runs of 1540 frames per condition, with batch size equal to 1 frame. Simulation was run with an NVIDIA© A30 GPU. Crosses indicate missing conditions.

### I. Biological plausibility

Here, we report on experimental data obtained from the literature, and evaluate the capacity of our simulator of fitting these empirical data. Using equations 7–10, our simulator accurately reproduces the relative phosphene brightness that was reported in the previous study for different stimulation amplitudes (*R*^2^ = 0.950, Figure 3). Figure 4 visualizes the effect of changing the stimulation parameters on the probability of phosphene perception, as estimated by our model. We compare our estimates with data reported by (6). As can be observed, our model accurately fits the reported effect of pulse width, frequency and train duration on the probability of phosphenes perception (*R*^2^ = 0.868).

**Fig. 3.**
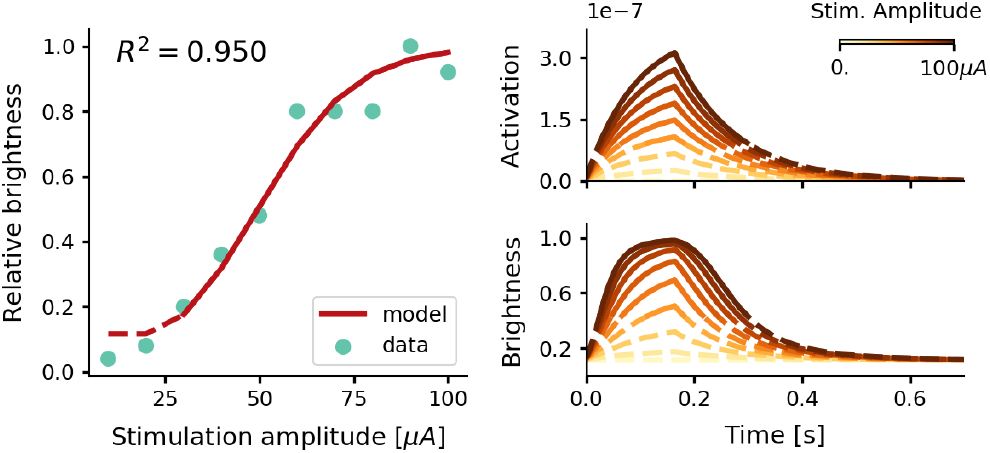
Estimate of the relative phosphene brightness for different stimulation amplitudes. The simulator was provided with a stimulation train of 166*ms* with a pulse width of 170*μs* at a frequency of 300*Hz* (see equations 7–10). Left: the predicted peak brightness levels reproduced by our model (red) and psychometric data reported by (6) (light blue). Note that for stimulation amplitudes of 20.0*μA* and lower, the simulator generated no phosphenes as the threshold for activation was not reached. Right: the modeled tissue activation and brightness response over time. Values below the 50% threshold for the tissue activation and the corresponding brightness values are displayed with dashed lines.

**Fig. 4.**
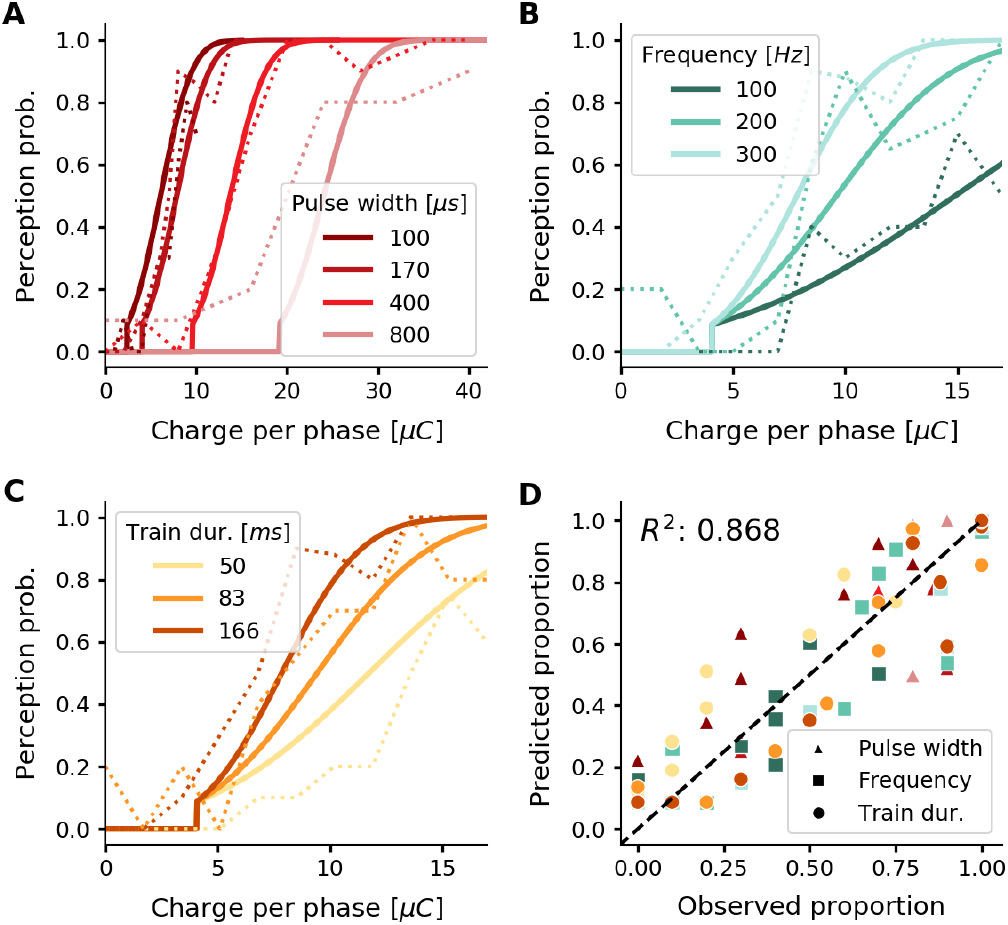
(a-c) Psychometric curves (solid lines) overlaid on experimental data (dashed lines) (ref. 6, Fig. 2a, b). The predicted probability of phosphene perception is visualized as a function of charge per phase for (a) different pulse widths, (b) different frequencies, and (c) different train durations. Note that rather than the total charge per trial, we report the charge per phase to facilitate easy comparison with aforementioned experimental data. In panel (d) the probabilities of phosphene perception reproduced with our model are compared to the detection probabilities reported in (ref. 6, Fig. 2a, b). Colors conform to the conditions in panels a, b and c

Figure 5 displays the results of exploring our simulator’s capacity of modeling temporal dynamics found in a previous published study by (23). For repeated stimulation at different timescales (intervals of 2 seconds, and intervals of 4 minutes), the measured brightness of a single phosphene is evaluated after fitting the memory trace parameters. The observed accommodation effects in our simulator are compared with the data from (23).

**Fig. 5.**
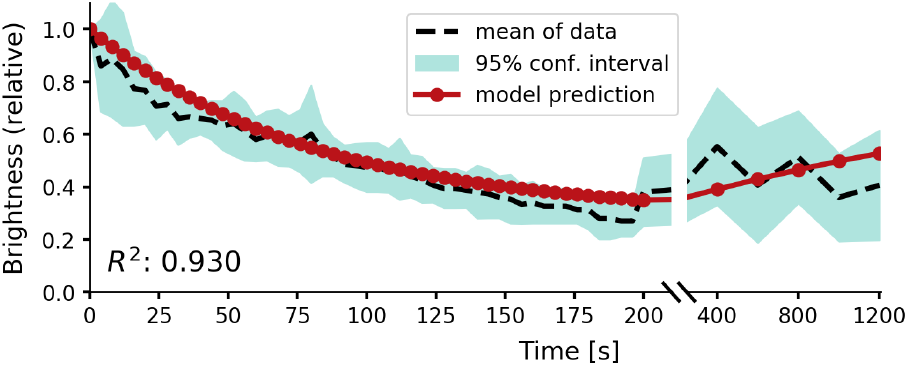
Relative brightness of a phosphene in response to repeated stimulation, overlaid on experimental results by (23). The stimulation sequence consisted of 50 pulse trains at a four-second stimulation interval, followed by five pulse trains at an interval of four minutes to test recovery. Please notice the split x-axis with variable scaling.

### J. Usability in a deep learning SPV pipeline

To validate that the simulator can conveniently be incorporated in a machine learning pipeline, we used our simulator in an existing SPV pipeline by (7) and performed several phosphene encoding optimization experiments, described below. In this pipeline, a convolutional neural network encoder is trained to process images or video frames and generate adequate electrode stimulation parameters. To train the encoder, a simulation of the prosthetic percept is generated by our differentiable phosphene simulator. This simulated percept is evaluated by a second convolutional neural network, the decoder, which decodes the simulated percept into a reconstruction of the original input image (Figure 6). The quality of the phosphene encoding is optimized by iteratively updating the network parameters of the encoder and decoder (simultaneously) using backpropagation of the reconstruction error. In addition to the reconstruction error, which measures similarity between the reconstruction and the input, we used a regularization term that measures similarity between the phosphenes and the input. For a more detailed description of the end-to-end optimization pipeline, see (7).

**Fig. 6.**
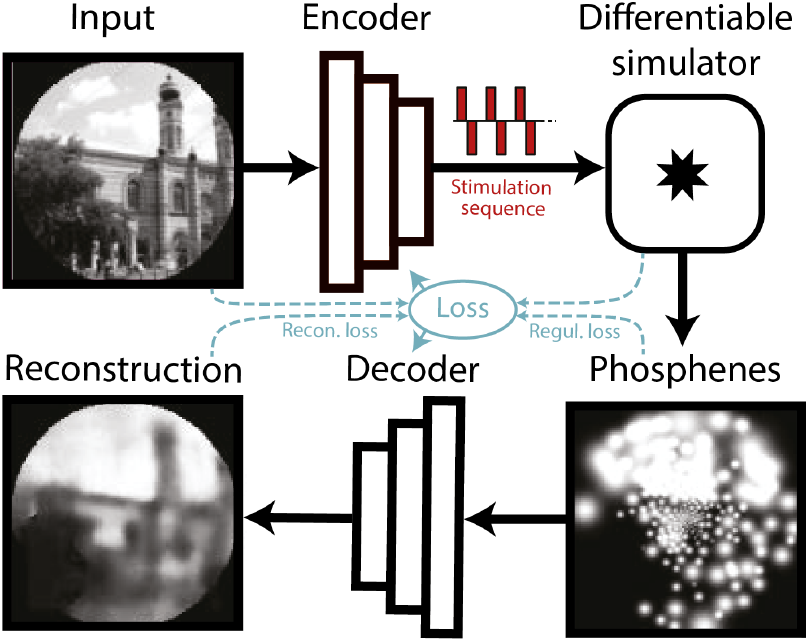
Schematic illustration of the end-to-end machine-learning pipeline adapted from (7). A convolutional neural network encoder is trained to convert input images or video frames into a suitable electrical stimulation protocol. In the training procedure, our simulator generates a simulation of the expected prosthetic percept, which is evaluated by a second convolutional neural network that decodes a reconstruction of the input image. The quality of the encoding is iteratively optimized by updating the network parameters using back-propagation. Different loss terms can be used to constrain the phosphene encoding, such as the reconstruction error between the reconstruction and the input, a regularization loss between the phosphenes and the input, or a supervised loss term between the reconstructions and some ground-truth labeled data (not depicted here). Note that the internal parameters of the simulator (e.g. the estimated tissue activation) can also be used as loss terms.

#### J.1. Dynamic end-to-end encoding of videos

In a further experiment, we explored the potential of using our simulator in a dynamic end-to-end encoding pipeline. We extended the previously published pipeline with 3D-convolutions (as an additional temporal dimension) to enable encoding of subsequent video frames. The model was trained on a basic video dataset with moving white digits on a black background (Moving MNIST Dataset, (58)). We used video sequences of five frames. The framerate of the simulation was set at five frames per second. We used a combination of two even-weighted mean squared error (MSE) loss functions: the MSE loss between reconstruction and input, and the MSE loss between the simulated phosphene representation and the input. Figure 7 displays several frames after training for 45 epochs (for a total of 810,000 training examples). We can observe that the model has successfully learned to represent the original input frames in phosphene vision over time, and the decoder is able to approximately reconstruct the original input.

**Fig. 7.**
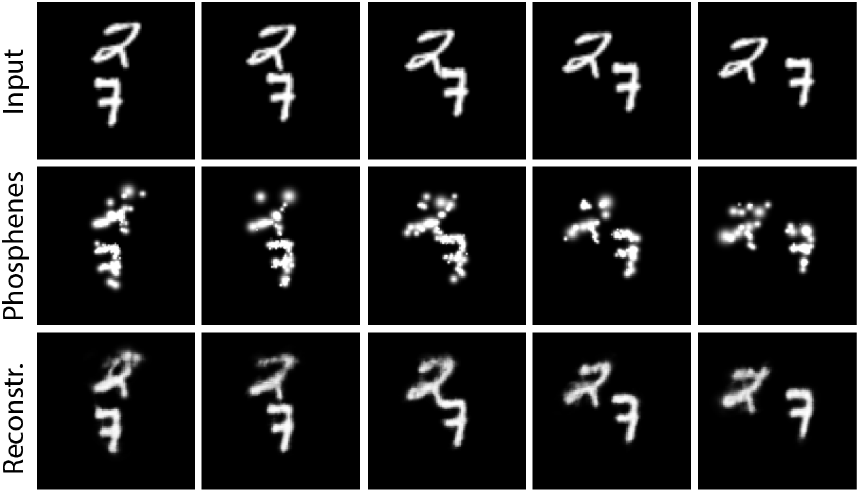
Results of training the end-to-end pipeline on video sequences from the moving MNIST dataset (58). Columns indicate different frames. Top row: the input frames; middle row: the simulated phosphene; bottom row: the decoded reconstruction of the input. This figure is best viewed in digital format.

#### J.2. Constrained stimulation and naturalistic scenes

In a second experiment, we trained the end-to-end model with a more challenging dataset containing complex images of naturalistic scenes (the ADE20K dataset (59)). In this experiment, we implemented the original pipeline described in (7) (experiment 4). The images were normalized and converted to grayscale, and we applied a circular mask such that the corners (outside the field covered by phosphenes) are ignored in the reconstruction task. The experiment consisted of three training runs, in which we tested different conditions: a free optimization condition, a constrained optimization condition, and a supervised boundary reconstruction condition. In the free optimization condition, the model was trained using an equally weighted combination of a MSE reconstruction loss between input and reconstruction, and a MSE regularization loss between the phosphenes and input images. After six epochs the model has learned to successfully find an optimal encoding strategy that can accurately represent the scene and allows the decoder to accurately reconstruct pixel intensities while qualitatively maintaining the image structure (see Figure 8). Importantly, note that the encoder has learned to encode brighter areas of the input picture by using large stimulation amplitudes (over 2000*μA*). The encoding strategy found in such an unconstrained optimization scheme is not feasible for real-life applications. In practice, the electrical stimulation protocol will need to satisfy safety bounds and it will need to comply with technical requirements and limitations of the stimulator hardware. For instance, rather than continuous stimulation intensities it is likely that the stimulator will allow for stimulation with only a number of (discrete) amplitudes. To evaluate whether our end-to-end pipeline can be harnessed to optimize the encoding in a constrained context, we performed a second training run (the constrained condition) where we reconfigured the encoder to output 10 discrete values between 0 and 128*μA*. We used straight-through estimation with a smooth staircase function to estimate the gradients during backpropagation. We increased the training stability by adapting the relative weights of the reconstruction loss and the regularization loss (to 0.999 and 0.0001, respectively). The results of the safety-constrained training after six epochs are visualized in Figure 8. Note that overall, the resulting phosphenes are less bright and smaller due to the lower stimulation amplitudes. Nevertheless, the decoder is able to accurately reconstruct the original input. One limitation is that we did not test the subjective interpretability for human observers. As not all information in the scene is equally important, it may be informative to further constrain the phosphene representation to encode specific task-relevant features. In a third training run (the supervised boundary condition) we validated whether our simulator can be used in a supervised machine learning pipeline for the reconstruction of specific target features, such as the boundaries between objects. Instead of using the input image as a reference, now the MSE is used between the reconstruction and a ground truth target image and between the phosphene representation and the target image. The ground truth semantic boundary targets were obtained by performing canny edge detection and subsequent line thickening on the semantic segmentation labels provided with the dataset. The results after training for 16 epochs are visualized in Figure 8. Note that the model successfully converged to a sparse phosphene encoding that selectively represents the object boundaries.

**Fig. 8.**
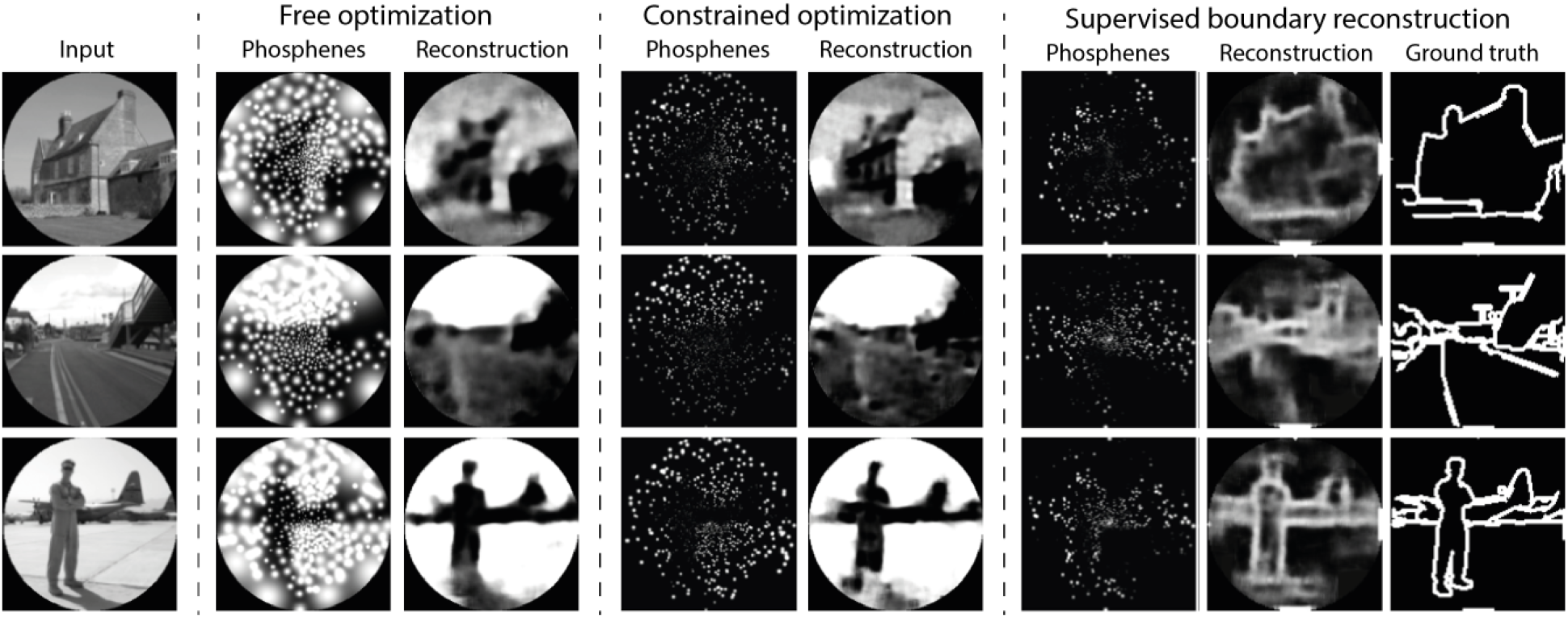
Results of training our simulator in an end-to-end pipeline on naturalistic images from the ADE20K dataset (59). In the constrained optimization condition and the supervised boundary reconstruction condition, the encoder was configured to output 10 discrete stimulation amplitudes within the safe range of stimulation (0 to 128*μA*). The selected images represent the first three categories in the validation dataset (’Abbey’, ‘Access Road’, ‘Airbase’). Note that the brightness is enhanced in the phosphene images of the constrained optimization and the supervised boundary condition by 40%. The figure is best viewed in digital format.

## Discussion

The aim of this study is to present a differentiable and biologically plausible phosphene simulator, which takes realistic ranges of stimulation parameters, and generates a phenomenologically accurate representation of phosphene vision. In order to achieve this, we have modeled and incorporated an extensive body of work regarding the psychophysics of phosphene perception. From the results presented in section I, we observe that our simulator is able to produce phosphene percepts that match the descriptions of phosphene vision that were gathered in basic and clinical visual neuroprosthetics studies that were conducted over the past decades. Running on a GPU, the simulator runs in real time, and as such, it could be used in clinical experiments with sighted volunteers. Furthermore, our proof-of-principle computational experiments, presented in section J demonstrate the suitability of our simulator for machine learning pipelines, aimed at improving the image processing and stimulation strategies. Here we discuss some implications of our findings.

### K. Validation experiments

#### K.1. Visuotopic mapping

The results presented in Figure 1 illustrate the value of including a visuotopic model based on spread of cortical activation to realistically estimate phosphene locations and size. Some previous studies have used a model of cortical magnification (34, 60) or visuotopic mapping (41, 42) in their phosphene simulations. However, our simulator is the first to incorporate empirical models of the current spread in cortical tissue (24, 25, 28) to simulate the effects of stimulation current on the phosphene size. The accurate modelling of this biophysical relationship can help to increase the validity of simulation studies and brings fundamental SPV research closer to addressing questions regarding the practical real-life requirements of a visual prosthesis. Furthermore, the explicit link between the modeled area of cortical activation and the simulated phosphene sizes and locations makes our software very suitable for including new receptive field modeling results. Collaborative international projects such as the PRIMatE Resource Exchange (PRIME-RE) offer advanced tools which allow to fit probabilistic retinotopic maps generated from large samples to any individual NHP brain (61, 62) and it is currently possible to accurately predict human cortical receptive field mapping based on anatomical scans (63, 64), or other retinotopic mapping strategies that do not rely on visual input (65, 66). This opens new doors for future research into the functionality of visual prostheses with limited visual field coverage. Thanks to the machine learning compatibility, the model can also be used for the pre-operative optimization of the implant placement and design (67).

#### K.2. Threshold and brightness

The results presented in Figures 3 and 4 indicate that our simulator closely models existing psychophysical data on the stimulation thresholds and phosphene brightness, for different electrical stimulation settings. Note that the effects found by (6) (that were modeled by us) are consistent with findings by other studies, which report brighter phosphenes for higher stimulation strengths (23), and a lower stimulation efficiency (i.e. higher total charge thresholds) for longer pulse trains or higher pulse widths and frequencies (20). Moreover, our results are in line with other computational models for detection thresholds in the somatosensory cortex (68, 69). Our results indicate how a leaky integrator model and normally-distributed activation thresholds, provide a suitable approximation of the tissue activation in cortical prostheses. Note that alternative, more complex, models can possibly predict the psychometric data more accurately. However, most probably, this will entail a trade-off with the simplicity and modularity found in our current simulator. Future research may further improve our understanding of the neural processing underlying the conscious perception of phosphenes, possibly borrowing insights from the domain of natural vision. More elaborate theories on this matter have been developed and tested in (70). More specific limitations and suggestions for future adaptations are discussed in section M.

#### K.3. Temporal dynamics

The results presented in Figure 5 reveal that the model accounts for experimental data on the accommodation in response to repeated stimulation in time periods up to 200 seconds. However, in contrast to the findings by (23), our simulator predicts a moderate recovery over the next 1000 seconds. Although we cannot provide an explanation for this difference, the modelled recovery largely stays within the 95% confidence interval of the experimental data. Similar to the other components of our simulator, the memory trace was chosen as a basic, yet effective, model of neural habituation. Possibly, a more complex, non-linear, model can more accurately fit the neurophysiological data, as discussed in section M. However, the presented model has the benefit of simplicity (there are only three parameters). Also, note that there is still some ambivalence in the clinical data. In contrast to the findings by (23), some studies have found accommodation over different time scales and sometimes no accommodation at all (6, 19, 22, 71). More research is required for a better understanding of the neural response after repeated or prolonged stimulation.

#### K.4. Phosphene shape and appearance

The appearance of phosphenes in our simulation (white, round, soft dots of light) are largely in line with previous reports on intracortical stimulation (6, 22, 23). However, more elongated shapes and more complex shapes have been reported as well (22, 24, 25). By using separately generated phosphene renderings, our simulator enables easy adjustments of the appearance of individual phosphenes. Additionally, we incorporated the possibility to change the default Gaussian blob appearance into Gabor patches with a specific frequency and orientation. Regarding the colour of phosphenes, there is still some ambivalence in reports, including descriptions of phospehene color ranging from black or white to different tones of color (23, 28, 72). Notably, increasing the stimulation amplitudes can lead the appearance to shift from colored to yellowish or white (23). This effect may be explained by the increased current spread for higher stimulation amplitudes, which is predicted to span multiple cortical columns coding for specific visual characteristics (e.g. orientation or colour), thus giving rise to phosphenes with amalgamated features (28). Currently, the limited amount of systematic data render it difficult to enable more accurate simulations of the variability in phosphene appearance.

### L. End-to-end optimization

#### L.1. Dynamic encoding

The results presented in Figure 7 demonstrate that our proposed realistic phosphene simulator is well-suited for the dynamic optimization in an end-to-end architecture. Our proof-of-principle video-encoding experiments are the first to explicitly optimize the stimulation across the temporal domain. This provides a basis for the further exploration of computationally-optimized dynamic stimulation patterns. Dynamic optimization of the stimulation may be necessary to counteract unwanted effects such as response fading due to accommodation after repeated or prolonged stimulation (23), or delayed phosphene perception after stimulation onset. The inclusion of a realistic simulator in the optimization pipeline enables researchers to exploit the optimal combination of stimulation parameters to obtain precise control over the required perception. Moreover, besides acquiring optimal control over the transfer function from stimulation to phosphenes, dynamic phosphene encoding could also prove useful to expand the encoded information along the temporal domain (21). Although this was not in the scope of the current study, our software is well-suited for simulation experiments that further investigate dynamic stimulation. Note that there remain some challenging perceptual considerations for the design of useful dynamics stimulation patterns (for an excellent review on asynchronous stimulation in relation to flicker fusion, form vision and apparent motion perception, please see (73)).

#### L.2. Constrained, efficient stimulation for natural stimuli

Our second optimization experiment addresses a more natural and realistic context. The results presented in Figure 8 demonstrate that our simulator is well-suited for the optimization of prosthetic vision to natural stimuli and that it can be configured to comply with constraints regarding the stimulation protocol. Note that the quality of the reconstructions for the constrained version of the encoder indicate that the model can still find an efficient information encoding strategy using a limited set of stimulation amplitudes (10 discrete values between 0 and 128μA). These results are in line with previous results on constrained end-to-end optimization, indicating that task-relevant information can be maximized under sparsity constraints (7). While in the current experiments the stimulation amplitude is maximized for the individual electrodes, future studies could investigate other sparsity constraints, such as a maximum total charge delivered per second across all electrodes. Ultimately, rather than an accurate overall portrayal of the visual surroundings, a visual prosthesis may need to prioritize task-relevant information. For this reason, in recent SPV research with sighted human observers much attention is devoted to semantic (boundary) segmentation for discriminating the important information from irrelevant background (10, 13, 48). Note that the explored image processing strategies in these behavioral studies are compatible with the automated optimization through an end-to-end machine learning pipeline. Our experiments exemplify how supervision targets obtained from semantic segmentation data can be adopted to promote task-relevant information in the phosphene representation. Furthermore, in addition to reconstruction of the input or labelled targets, another recent study experimented with different decoding tasks, including more interactive, goal-driven tasks in virtual game environments (17). Although these proof-of-principle results remain to be translated to real-world tasks and environments, they provide a valuable basis for further exploration. Ultimately, the development of task-relevant scene-processing algorithms will likely benefit both from computational optimization experiments as well as exploratory SPV studies with human observers.

#### L.3. Interpretability and perceptual correspondence

Besides the encoding efficiency (characterized by the computational decodability of task-relevant information), it is important to consider the subjective interpretablitity of the simulated phosphene representation. From the results in Figure 8 it can be observed that the model has successfully learned to preserve correspondence between the phosphene representation and the input image in all of the training conditions. However, as a more formal analysis was outside the scope of this study, we do not further quantify the subjective interpretability. In our model the subjective interpretability was promoted through the regularization loss between the simulated phosphenes and the input image. Similarly, a recent study adapted an auto-encoder architecture designed to directly maximize the perceptual correspondence between a target representation and the simulated percept in a retinal prosthesis, using basic stimuli (12). The preservation of subjective interpretability in an automated optimization pipeline remains a non-trivial challenge, especially when using natural stimuli. This may hold even more for cortical prostheses, as the distinct punctuate phosphenes are in nature very dissimilar from natural images, possibly hampering perceptual similarity metrics that rely on low-level feature correspondence. Regardless of the implementation, it will always be important to include human observers in the optimization cycle to ensure subjective interpretability for the end user (74, 75).

### M. General limitations and future directions

#### M.1. Performance and hardware

There are some remaining practical limitations and challenges for future research. We identify three considerations for future research related to the the performance of our model and the required hardware for implementation in an experimental setup. Firstly, although our model runs in realtime and is faster than state-of-the art simulation for retinal prostheses (15), there is a trade-off between speed and the memory demand. Therefore, for higher resolutions and larger number of phosphenes, future experimental research may need to adapt a simplified version of our model - although most of the simulation conditions can be run easily with common graphical cards. Secondly, a suggestion for follow-up research, is to combine our simulator with the latest developments in mixed reality (XR) to enable immersive simulation in virtual environments. More specifically, a convenient direction would be the implementation of our simulator using the Cg shader programming language for graphics processing, which is used in 3D game engines like Unreal Engine, or Unity 3D, as previously demonstrated for epiretinal simulations by (38). Thirdly, and lastly, future studies could integrate our simulation software with eye-tracking technology. Due to the retinotopic organization of visual cortex, cortical stimulation leads to phosphenes that are not fixed to a position in space, but change after every eye movement. These effects should be taken into account for a faithful simulation of the experience of a prosthesis user. Furthermore, as has been demonstrated in previous SPV studies (34, 35), one could investigate the potential benefits of including an eye tracker in the prosthetic hardware to allow for sampling of the visual environment using eye movements. Note that testing such gaze-assisted processing does not require any changes to our simulator. It merely involves processing the gaze-centered image as opposed to the entire camera input.

#### M.2. Complexity and realism of the simulation

There are some remaining challenges regarding the realistic simulation of the effects of neural stimulation. A complicating factor is that cortical neuroprostheses are still in the early stages of development. Neurostimulation hardware and stimulation protocols are continuously being improved (21), and clinical trials with cortical visual neuroprostheses are often limited to small numbers of blind volunteers (11, 76). Therefore, it is no surprise that the amount of data that is available at the present moment is limited, often open for multiple interpretations, and sometimes contains apparent contradictory information. Notably, the trade-off between model complexity and accurate psychophysical predictions is a recurrent theme in the validation of the components implemented in our simulator. These factors play a role in some of the potential limitations of our current simulator. Here we name a few of the important limitations and some interesting directions for future research. Firstly, in our simulator, phosphenes are only rendered when the activation is above threshold and vice-versa. This might be an inaccurate depiction of the perceptual experience of an implant user, and in reality the distinction may be less strict. The conscious perception of phosphenes requires considerable training and the detection process is influenced by attention (6). Although our implementation is effective for modeling the psychometric data, alternative implementations could also be considered. The perceptual effect of different simulated phosphene threshold implementations for sighted subjects remains to be evaluated in future SPV work. Secondly, the leaky integrator and the memory trace that are implemented in our simulator might be an oversimplified model of tissue activation in the visual cortex. This means that some non-linear dynamics might be missed. Also, several studies reported that higher stimulation amplitudes may give rise to double phosphenes (18, 19, 23, 77), or a reduction of phosphene brightness (23). Furthermore, interactions between simultaneous stimulation of multiple electrodes can have an effect on the phosphene size and sometimes lead to unexpected percepts (6). Further clinical data could help to improve our understanding of these non-linear dynamics. A third limitation is that our simulator currently only models responses of V1 stimulation. Future studies could explore the possible extension of modeling micro-stimulation of higher visual areas, such as V2, V3, V4 or inferotemporal cortex. In previous NHP research, reliable phosphene thresholds could be obtained with the stimulation of in V1, V2, V3A, MT (78). Furthermore, IT stimulation has shown to bias face perception (79). Similar effects have been confirmed in human subjects, and previous work has demonstrated that electrical stimulation of higher order visual areas can elicit a range of feature-specific percepts (80–82). Our simulator could be extended with maps of higher cortical areas with clear retinotopy, and an interesting direction for future research will be the implementation of feature-specific percepts, including texture, shape and colour.

### N. Conclusion

We present a biologically plausible simulator of phosphene vision. This simulator models pshychophysical and neurophysiological findings in a wide array of experimental results. Its phenomenologically accurate simulations allow for the optimisation of visual cortical prosthesis in a manner that drastically narrows the gap between simulation and reality, compared to previous studies of simulated phosphene vision. It can operate in real time, therefore being a viable option for behavioural experiments with sighted volunteers. Additionally, the PyTorch implementation and its differentiable nature makes it a good choice for machine learning approaches to study and optimize phosphene vision. The modular design of our simulator allows for straightforward adaptation of novel insights and improved models of cortical activation. In summary, our open-source, fully differentiable, biologically plausible phosphene simulator aims to provide an accessible bedrock software platform that fits the needs of fundamental, clinical and computational vision scientists working on cortical neuroprosthetic vision. With this work, we aspire to contribute to increasing the field’s translational impact.

## ACKNOWLEDGEMENTS

This work was supported by two grants (NESTOR, INTENSE) of the Dutch Organization for Scientific Research (NWO) and the European Union’s Horizon 2020 research and innovation programme (under grant agreement No 899287). We thank Xing Chen for help with the compilation and reviewing of relevant literature regarding phosphene perception.

## Footnotes

1 The source code of our simulator can be retrieved from Github: https://github.com/neuralcodinglab/dynaphos. The latest stable release can be installed using pip: `$ pip install dynaphos`

2 The example video with the simulated phosphene output can be downloaded via [this link](https://github.com/neuralcodinglab/dynaphos/blob/main/examples/example_video.mp4).

